# Genetic algorithm-based personalized models of human cardiac action potential

**DOI:** 10.1101/712406

**Authors:** D.N. Smirnov, R.A. Syunyaev, R.M. Deviatiiarov, O.A. Gusev, K.K. Aras, A.C. Koppel, I.R. Efimov

**Affiliations:** Moscow Institute of Physics and Technology, Dolgoprudny, Russia; Kazan Federal University, Kazan, Russia; George Washinton University, Washington, USA; Sechenov University, Moscow, Russia

## Abstract

We present a novel genetic algorithm-based solution to determine the set of cardiomyocyte model parameters based on experimental human action potential (AP) recordings. The novel approach is based on AP waveform dependence on the heart rate. In order to find the steady-state solution, optimized parameters include conductivities of ionic channels and exchangers augmented by slow variables, intracellular sodium concentration and sarcoplasmic reticulum calcium load. The algorithm is enhanced by a novel mutation operator, based on Cauchy distribution along a random direction in parameter space. We also demonstrate that increasing the number of elite organisms up to 7% results in faster convergence. Test runs indicate that algorithm error is below 5% for I_Kr_, 7% for I_K1_ and I_Na_ and 13% for I_CaL_. Experimental signal-to-noise ratio above 28 dB was sufficient for high quality algorithm performance. The algorithm validation using optical mapping recordings of human ventricular AP demonstrated low error that was below 6 mV for AP waveform and less than 16 ms for AP duration. Further validation of the personalized models was done using mRNA expression profile of two donor hearts. The mRNA-based model reproduced AP waveform dependence on cycle length with 13 mV accuracy and AP duration with 20 ms accuracy.

## Introduction

Over the past few decades, mathematical models of cardiac electrophysiology came a long way in terms of complexity and the area of application. Recent advances in computational cardiac electrophysiology make clinical application of computer models possible due to personalization of tissue geometry and fibers orientation^1^. However, despite these advances, single cell-mathematical models remain non-personalized, being built from data collected in many laboratories from different subjects. While tissue-specific, person-specific and pathology-specific gene expression profiles affect AP propagation, these differences are usually not accounted for: tissue level simulations are usually based upon the same average elements of the model.

A number of publications utilized genetic algorithms (GA) to find a set of cell model parameters reproducing experimental AP^2–5^. GA apply evolutionary principles to computational models aiming to find the best solution fitting experimental data. Initially, a number of model “*organisms*” with random parameter values is generated. After that, “*selection*” operator is applied to the first *generation* of models, passing the models with higher values of *fitness function* to the “*mating pool*”. Usually, fitness function is based on Euclidian distance: squared difference between model and experiment. *Mutation* and *crossover* operators are then applied to the models in the mating pool. The former modifies model parameters according to some probability distribution function. In the simplest GA setting the crossover operator exchanges the parameter values between organisms with fixed probability, however more complicated modifications of crossover operators, such as Simulated Binary Crossover were shown to improve algorithm performance^6^. Modified models are then passed to the next generation, new fitness function values are calculated, and the same set of genetic operators is applied iteratively until desirable goodness of fit is reached.

An obvious advantage of GA is that these algorithms are *perfectly parallel* problems, making its parallel computation implementation straightforward: the slowest part of the algorithm, which is fitness function calculation, could be performed independently for different organisms, while communication between tasks is limited to a small array of parameters. However, effective implementation of GA requires modification of genetic operators for each particular set of problems (this fact is often referred to as “*no free lunch theorem*” in the literature^7^). One of the goals of the current study was to develop GA implementation suitable for cardiac electrophysiology models.

Another well-known limitation of optimization algorithm applied to electrophysiological models is the absence of a unique solution. As was noted previously^8^ same AP waveforms could be reproduced by computer models with different sets of parameters. Recently an approach combining stochastic pacing and complicated voltage-clamp protocols were proposed to overcome the problem^4,8^. However, these approaches are limited to single cell voltage-clamp recordings, which are not feasible in clinical electrophysiology. The aim of the current study was to develop a technique that would allow finding the unique solution using cardiac tissue AP recordings by optical mapping, microelectrode, or monophasic recordings. One possible approach to achieve this goal is to utilize so-called restitution property, which is AP dependence on heart rate or pacing cycle length (PCL). Restitution property allowed us to substantially narrow down the range of output model parameters.

## Methods

### Computer simulations

The human ventricular cell electrophysiology was simulated with O’Hara-Rudy model^9^. The genetic algorithm (GA) is based on Bot et al.^3^ with the following important modifications (green-tinted boxes on Fig 1A):

**Figure 1.**
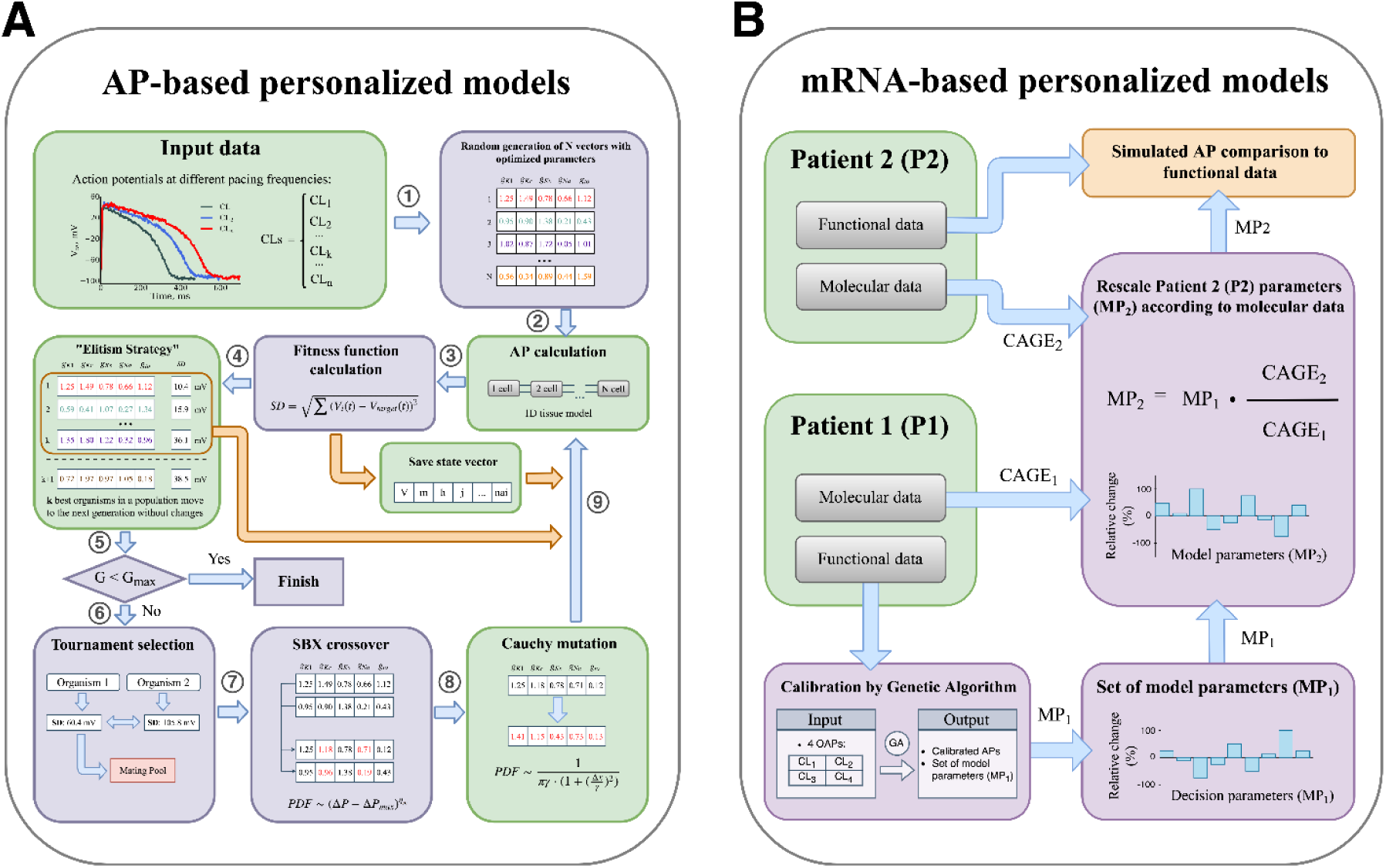
Genetic Algorithm (GA) and CAGE-based personalization block diagrams. (**A**) Genetic algorithm schematic diagram. At the initial stage a set of randomly generated organisms is created, each of which is determined by a vector of scaling factors for optimized model parameters (step 1). For each organism AP waveforms are calculated at several pacing frequencies and compared with the input APs (steps 3, 4). Organisms with the lowest SD value are saved and directly copied into the subsequent generation, replacing the worst organisms (orange arrow). State vectors (intracellular concentration, gating variables etc) are also saved after each short simulation and reused as initial state during simulation in the next generation. After selection (step 6) the fittest organisms form the mating pool and modified by SBX crossover and Cauchy mutation (step 7, 8). Modified organisms move to the next generation (9). Process of AP and fitness function calculation, selection, crossover and mutation and elite replacement is repeated until the end of generations limit. The algorithm is based on ^3^, modifications of the original algorithm are highlighted by green. (**B**) Algorithm verification with molecular (mRNA expression) and functional data (optical mapping). Patient 2 model parameters (MP_2_) were determined by GA. Patient 1 parameters (MP_1_) were rescaled proportional to Patient 2 mRNA expression level (CAGE_2_) and verified against functional data.

1. **Input data** is steady-state AP waveforms recorded at several PCL. The algorithm is optimized to use optical mapping recordings, thus absolute transmembrane potential (TP) values are not known. Conventional least-squares technique is used to find scaling and shift coefficients of AP waveform: *i.e.* experimental AP is rescaled and shifted to minimize deviation between the experiment and the model.
2. **AP calculations.** Intercellular interactions affect AP waveform. On the other hand, our simulations have shown that within the physiological range of conduction velocities (CV), 20-100 cm/s, exact gap junction conductivity value does not affect AP (S1B Fig). Therefore, during a GA run each model was simulated as 1D-tissue with 5 mS/μF conductivity between cells resulting in CV of 27 cm/s. The 1D-string of cells was 30 cells long, since we have found it enough to exclude boundary effects on the central cell in case of CV slower than 27 cm/s (S1A Fig).
3. **Fitness function.** We used standard deviation formula to evaluate how close is a given organism AP to input data at particular PCL:

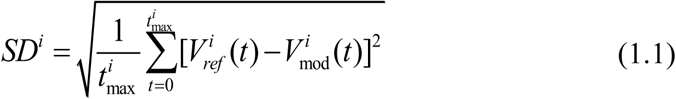

where 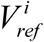 is a baseline TP, 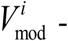 is a TP of simulated AP, 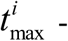 time length of input data recording. The fitness function was calculated as a weighted sum of *SD*^*i*^ corresponding to different PCLs:

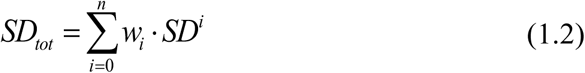

where *w*_*i*_ corresponds to a weight coefficient. Weights were taken equal for all PCLs unless otherwise noted.
4. **Save state vector.** Since reaching steady state for every organism during a GA run is computationally impossible, each AP is saved after a short simulation. We used 9 stimulations before fitness function evaluation, since we have found odd number of stimulations helpful to exclude possible 1:1 alternans in the output model. After that state vector (ionic concentrations, gating variables *etc.*) are saved for each organism. These state vectors are used as initial state in the next GA generation. This approach allows modifying parameters and reaching steady state at the same time.
5. **“Elitism strategy”.** Since genetic operators tend to spoil a “good” solution, good organism is passed to the next generation without any changes replacing the worst^10^. We have found that increasing the number of elite organisms to about 7% of the whole population is optimal.
6. **Cauchy mutation.**
  A. We have found that original polynomial mutation tends to trap solution in the local minima, which is a well-known problem of optimization algorithms. This is especially important in terms of intracellular ionic concentrations; therefore, we modified the mutation operator using Cauchy distribution:

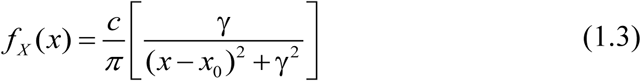

where *f*_*X*_ is a probability density of the distribution; *x*_0_ corresponds to unmutated parameter value; *γ* = 0.18* *x*_*O*_’_ *Hara*–*Rudy*_ is the half-width of the half-maximum of the distribution; 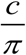 is a normalization constant resulting from a limited range of parameter values, varied between 0.01· *x*_*O*’*Hara*–*Rudy*_ and 4.0· *x*_*O*’*Hara*–*Rudy*_; *x*_*O*’*Hara*–*Rudy*_ is original O’Hara-Rudy model parameter value
  B. Usually mutation operator is applied to each parameter separately with some fixed probability^11^, we have found that it has adverse effects on algorithm convergence. Instead, we choose random direction in multi-dimensional parameter-space and mutate the parameter vector in this direction. See more details in the “Results” section.
  C. The slowest variables (intracellular Na^+^ and network sarcoplasmic reticulum Ca^2+^ concentrations) are included in parameters vector and mutated as usual model parameters. This approach allows algorithm to reach the model steady state much faster. Note that intracellular concentrations are different for different PCL, consequently, there are separate values for each pacing frequency.

In order to verify the algorithm precision, simulated APs with an arbitrary set of model parameters were used as input data. The input model was paced until reaching steady state (1000 s) at several PCLs: 217 ms, 225 ms, 250 ms, 300 ms, 500 ms, 1000 ms, 2000 ms, unless noted otherwise.

### Multidimensional data visualization

A. **Andrews curves** were used to visualize similarity between organisms within a single generation. Each organism is represented as 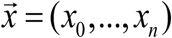, where *x*_*i*_ (*i* =1…*m*) are GA optimized parameters. Each point of this vector defines a Fourier series^12^:

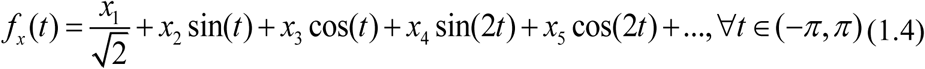

Thus, every organism from a generation is depicted as a single curve. An important feature of Andrews curves is a linear mapping of the distance between organisms in Euclidean metrics: if two curves stay close to each other for all ^*t*^ values, then corresponding organisms are close in parametric space as well.
B. **Principal Component Analysis** technique was used to visualize convergence in multidimensional parametric space. Organisms parameters of *p* compared generations form a matrix *X* of size *m*×*n*\* *p*, where *m* is the number of organisms and *n* is the number of optimized parameters. According to the principal component method, matrix *X* decomposed into a multiplication of two matrices *T* (scores matrix) and *P* (loading matrix) plus residual matrix *E* :

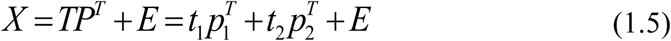

where *p*_*i*_, (*i* =1, 2) correspond to loading matrix rows, columns *t*_*i*_, (*i* =1, 2) of matrix *T* are Principal Components (PC) being the organisms parameters in the new coordinate system. *P* is a transformation matrix from initial variables space to 2D space of principal components.
C. In order to estimate organism parameters convergence in the principal components space we used **Mean Cluster Error** (MCE) and **Standard distance** (SDist) metrics (Fig. 7A):

a. Mean Cluster Error represent the distance between *R*(*x*_0_, *y*_0_) point corresponding to precise solution (input model value) and *A*(*x*_*c*_, *y*_*c*_) corresponding to cluster geometric center calculated within a single generation:

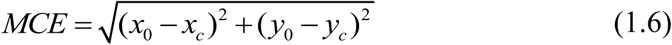
b. Standard Distance was used to estimate cluster size:

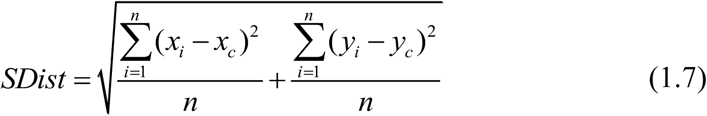

where *x*_*i*_ and *y*_*i*_ are principal components of a given organism, (*x*_*c*_, *y*_*c*_) is the cluster geometric center and *n* is the total number of organisms within a generation.

### Donor Heart Procurement

All studies using human heart tissue were approved by the Institutional Review Board (Office of Human Research) at the George Washington University. In total, for this study, we procured from Washington Regional Transplant Community in Washington, DC discarded ventricular tissues from 7 deidentified donor human hearts, which were unsuitable for transplantation. All hearts were arrested using ice-cold cardioplegic solution in the operating room and tissue was transported to the laboratory for dissection and electrophysiological experiments.

For subsequent mRNA analysis left ventricular tissue samples were dissected, submerged in RNAlater (Invitrogen) for 24 hours at 4°C, and stored at −80° C until the extraction of RNA. Total RNA was extracted from these samples using the RNeasy Fibrous Tissue Mini Kit (Qiagen) according to the manufacturer’s instructions, from approximately 30 mg samples. At the end of the extraction, spin columns were eluted with the eluate to increase the RNA yield. Total RNA concentration and purity were determined on an Eppendorf BioPhotometer D30. Acceptable purity, as quantified by both the 260/280 and 260/230 absorbance ratios, was between 1.8 and 2.2. It was also evaluated for integrity by electrophoresis on 1.2% agarose gels with 0.25 μg/ml ethidium bromide.

### Optical Mapping

Human left ventricular wedge preparations were used for experiments, as described previously^13^. Briefly, wedges from the posterolateral LV free wall perfused via the left marginal artery were dissected, cannulated, and mounted in a tissue chamber with 4 surfaces (epicardium, endocardium, and the 2 transmural sides) facing 4 CMOS cameras of the optical apparatus^13^. Preparations were perfused with oxygenated Tyrode solution maintained at 37°C, with a perfusion pressure of 60 to 80 mm Hg. The preparation was washed with 2 L of Tyrode solution to remove excess transplant solution and restore basal electrophysiology. Tissue was immobilized by blebbistatin (10–15 μM) to suppress motion artifacts in optical recordings, without adverse electrophysiological effects^14^. Di-4-ANBDQBS was used to map transmembrane potential as described previously^15^. Platinum-iridium tipped bipolar pacing electrode was placed on the epicardial surface. Optical action potentials were mapped from ≈5×5 cm field of view from the 4 surfaces using 4 MiCAM05 (SciMedia, CA) CMOS cameras with high spatial and temporal resolutions (100×100 pixels; sampling frequency, 1 KHz).

### Cap-Analysis of Gene Expression (CAGE)

This study was done on RNA extracted from human tissues procured as described above. After total RNA was extracted, as previously described, 5 μg of total RNA (260/280 2.0±0.01, 260/230 1.9±0.15, RIN 8.0±0.66) were submitted for 5’nAnT-iCAGE libraries preparation according to standard protocol^16^, sequenced and demultiplexed on Illumina HiSeq2500 High throughput mode (50nt single end). Libraries *in silico* processing performed in Moirai system^17^. This protocol includes quality control (fastx_trimmer: −Q33, −l 47), N base, and rRNA trimming (rRNAdust v1.0) with subsequent alignment to the human genome version hg19 through Burrows Wheeler Aligner (BWA). The median mapping ratio was 0.88±0.02 and median depth of 11.7M. CTSS (CAGE transcription start sites) and clusters of CAGE signal generated by applying python scripts: level1 and level2^18^ on BAM files resulted in median 1.27M of CTSS and 13.1K of putative promoter regions, where ~12.1K overlap promoters from FANTOM5^19^. Finally, 11612 predicted promoters were associated with 10355 genes through RefSeq and Ensembl transcripts obtained from UCSC^20^ by extending the searching area of its 5’ ends in ±500bp. CAGE promoters for the key genes were manually curated by visualization in Zenbu browser^21^. TPM (tags per million) normalized CAGE counts were submitted to edgeR package for R for differential expression analysis according to the protocol^22^.

### Algorithm verification with experimental data

In two donor hearts after collecting tissue samples for CAGE, ventricular wedge preparations were made for optical mapping as described above. Optical APs were recorded from endocardial surface of the preparation while the tissue was paced epicardially until reaching steady state (100 s) at 4 different PCLs: 2000 ms, 1000 ms, 500 ms and 300 ms. We observed some variation in AP waveform over endocardial surface, therefore a single pixel recording with higher APD, upstroke velocity and overall signal-to-noise ratio was chosen manually and used as input data for GA. Low-pass filter was not used, to avoid AP waveform distortion. 60-Hz hum was removed with narrow band stop IIR Butterworth filter. Ensemble averaging over a series of APs recorded from the same pixel was used to further improve signal to noise ratio (SNR).

When GA was applied to experimental recording, we could not directly verify the precision of GA output ionic channel conductivities. Instead, the following indirect approach was used (Fig 1B). *Patient 2* functional data from optical mapping was used as input for GA. Output model parameters (ionic channels conductivities) were rescaled proportional to differences in mRNA expression between two patients (similar approach was used in ^9^). We considered differences in SCN5A, KCNH2, KCNJ2, KCNQ1, CACNA1C, KCNA4, ATP1A1, SLC8A1, ATP2B4, RYR2, ATP2A2, CALM1, CAMK2D, CALM1 to be represented in the model by INa, IKr, IK1, IKs, ICaL, Ito, INaK, INCX, IpCa, Jrel, Jup, CMDN, CaMKII correspondingly. Differences in these genes level of expression among 7 patients’ hearts are shown in Fig 2, *Patients 1* and *2* are highlighted by orange and grey colors. The resulting *Patient 1* model was compared to *Patient 1* functional data as described below in “Results” section.

**Figure 2.**
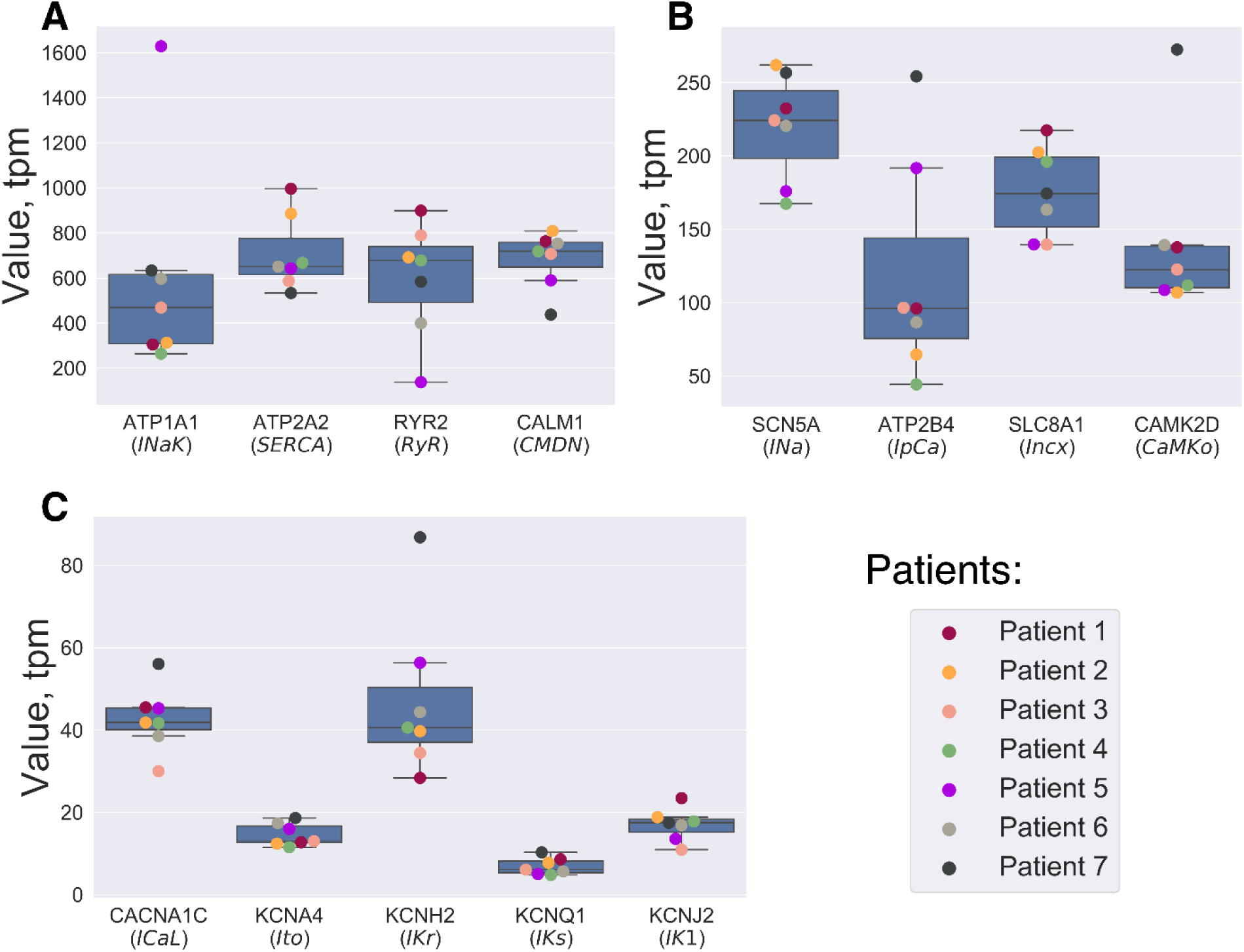
The mRNA expression data measured via Cap Analysis of Gene Expression (CAGE). (**A-C**) The mRNA expression level measured in 7 donor hearts. Only genes used for rescaling model parameters are shown. Outliers were determined by IQR method.

## Results

### Intracellular concentrations

The immediate consequence of using steady-state AP waveform dependence on PCL as GA input data is that output model AP should be steady state as well. The direct approach to the problem is to pace every organism for a long time during a GA run. However, this solution is computationally very expensive: successful GA convergence requires at least 100 organisms and 100 generations^2,4^. On the other hand, arbitrary initial state of the model requires at least 100 s to stabilize intracellular ionic concentrations. These slow transients are mostly determined by intracellular Na^+^ concentration and Ca^2+^ sarcoplasmic reticulum load changes. Moreover, given different initial state, intracellular concentration converges to different values as exemplified in Fig 3B-C. Fig 3A shows that these differences may have significant effects on the steady-state AP waveform: resting membrane potential (RMP) difference is 4.8 mV, AP duration difference is 15 ms.

**Figure 3.**
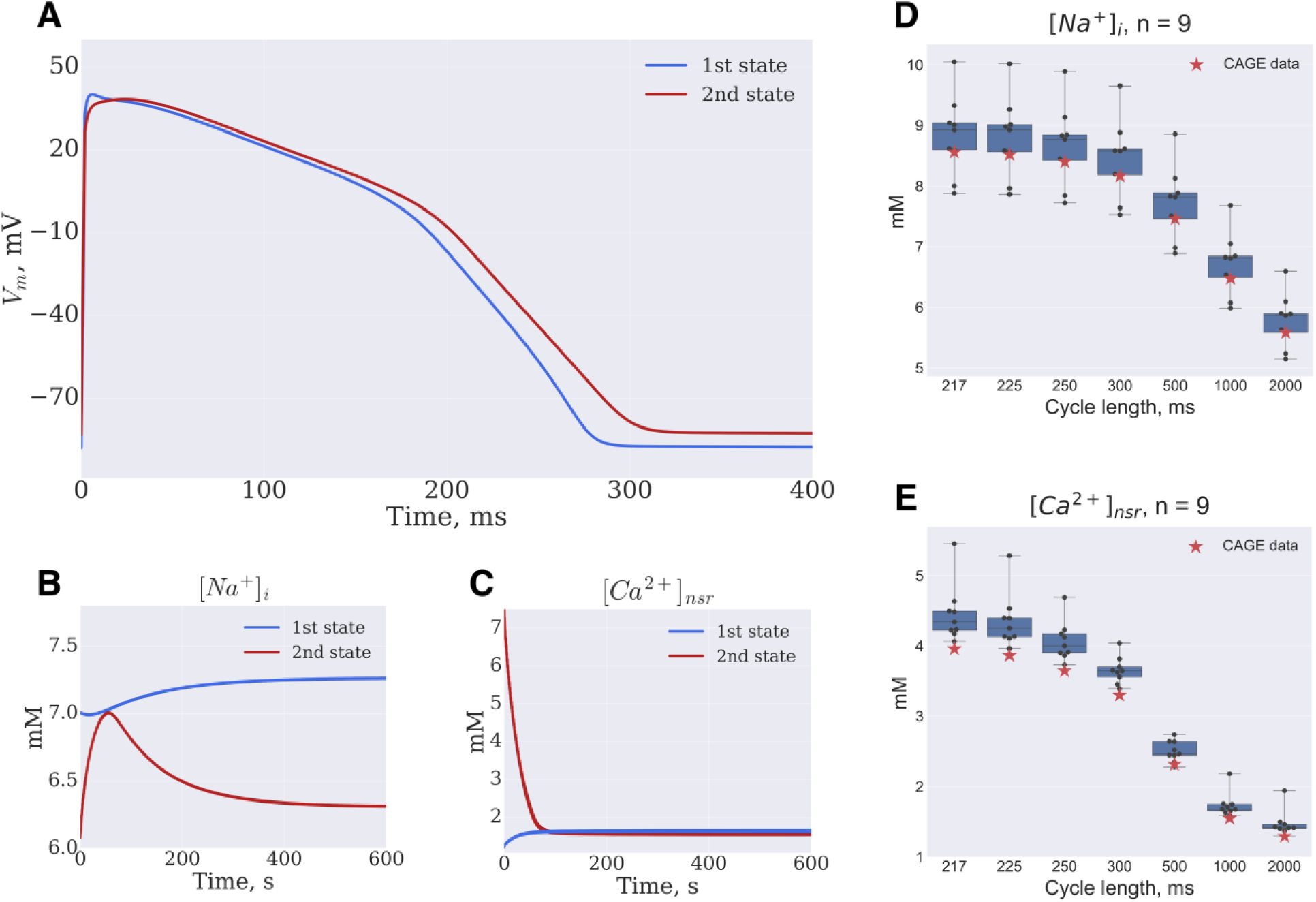
Saving organisms state vectors. (**A**) The single-cell model was paced at 1Hz frequency for 600 s, from different initial [Na^+^]_i_ and [Ca^2+^]_nsr_ concentrations, resulting in different steady-state resting potential and APD. (**B, C**) [Na^+^]_i_ and [Ca^2+^]_nsr_ concentrations changes during the simulations. (**D, E**) Box-and-whiskers plots depict [Na^+^]_i_ and [Ca^2+^]_nsr_ dependence on PCL as determined by 9 GA runs. Red stars are input model values.

Instead of the direct approach, we have evaluated the fitness function after a short run (9 stimulations at each PCL). The state variables at each PCL are saved and reused as initial state at the next generation (Fig 1A). In order to further speed-up intracellular concentrations convergence, Na^+^_i_ and Ca^2+^_nsr_ for each PCL were added to parameters vector and were susceptible to mutation and crossover operators. The resulting algorithm allowed us to determine the steady-state concentration with 1.5 mM precision after 700 generations (Fig 3D-E).

### Mutation operator

Usual GA approach is to treat parameters separately by mutation operator, *i.e.* there is a fixed probability to mutate each parameter^11^. As demonstrated in Fig 4B this approach (“*point mutation*”) results in a low probability to modify several parameters at the same time. SD dynamics averaged on 9 GA runs (Fig 4C) demonstrate that coordinate descent resulting from *“point mutation”* halts algorithm convergence after 100 of generations. Therefore, in our GA implementation random direction in parameter space is chosen by mutation operation and the whole set of parameters is modified at the same time (“*vector mutation*”, Fig 4A) resulting in better convergence (Fig 4C). Fig 4D shows that after 700 generations *vector mutation* estimates parameters much better than *point mutation:* for example, the error is 7% vs 20% for I_K1_, 5% vs 9% for I_Kr_, 7% for 24% for I_Na_.

**Figure 4.**
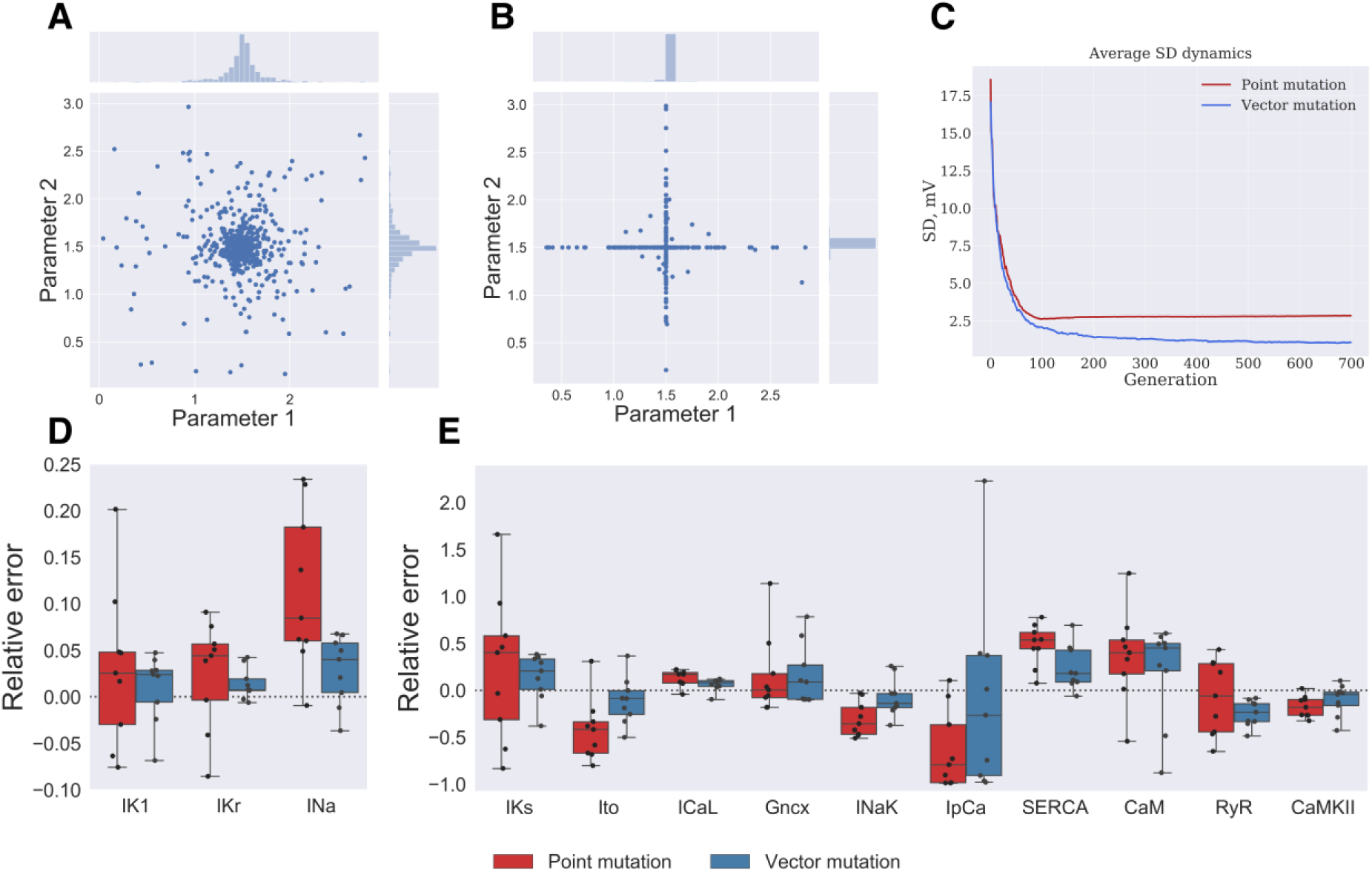
Random mutation direction in multi-dimensional parameter space. **(A)** Vector mutation: random mutation direction choice, blue points indicate parameter values after mutation. **(B)** Point mutation: each parameter value is mutated with fixed probability, which results in low probability to mutate parameter value in diagonal direction. (**C**) SD dynamics averaged over all organisms of 9 GA runs plotted against generation number. Red and blue lines correspond to point and vector mutations, respectively. (**D, E**) Objective parameters distribution for 9 GA runs with point mutation (red boxes) and vector mutation (blue boxes). Dashed line corresponds to the input model parameter values.

As mentioned above (see “Methods” section and “Intracellular concentrations” paragraph) Na^+^_i_ and Ca^2+^_nsr_ variables were included in the mutation parameters vector. Although the best organism is considered “elite” and unaffected by mutation operator, its AP and intracellular concentrations are still changing because of the model convergence to the steady state. These changes result in the best organism fitness function decrease (SD increase). On the other hand, crossover operator results in a number of organisms close to the best one in parametric space, *i.e.* GA have a tendency to converge to the local minimum: as shown by Andrews curves in Fig 5C the whole population is in the immediate vicinity of a single solution. Consequently, the best organism is constantly being replaced by one with similar parameters (and concentrations) values and model steady state might be never reached. Fig 5D-E (red lines) exemplifies this case: best organism (organism with the lowest SD) intracellular concentrations are constantly fluctuating because of the best organism replacement, and Ca^2+^_nsr_ don’t converge to the proper value (dashed line). To address the problem, in our GA implementation mutated parameters follow Cauchy distribution instead of the usual polynomial distribution (Fig 5A). As shown in Fig 5B, Cauchy mutation results in a wider variation in a population overlapping with input model values. Moreover, the best organism is unchanged for a number of generations allowing it to reach steady state (Fig 5D-E; after generation №500). Fig S2 illustrates the best organism intracellular concentrations averaged over 9 GA runs, showing that Cauchy mutation results in better convergence then polynomial.

**Figure 5.**
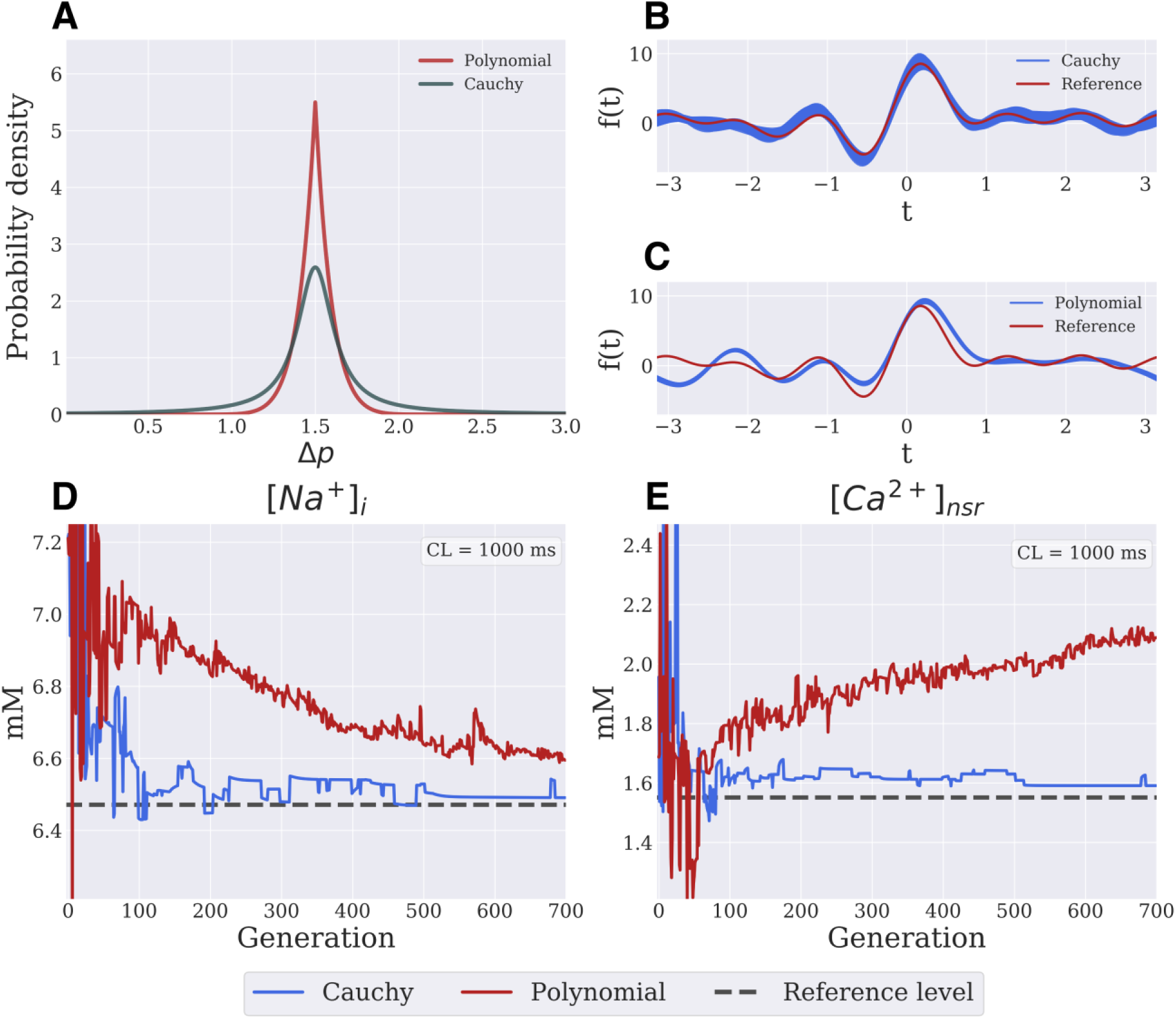
Cauchy mutation. (**A**) Cauchy (dark green line) and polynomial (red line) distributions comparison. (**B**,**C**) Cauchy distribution results in a broader range of parameters search than polynomial distribution as demonstrated by superimposed Andrews curves for all organisms on generation № 700 of a typical GA run (blue lines). Red line corresponds to input model set of parameters in both plots. (**D**,**E**) Best organism intracellular [Na^+^]_i_ and [Ca^2+^]_nsr_ concentration plotted against generation number in a sample GA run. Cauchy mutation (blue) allows the algorithm to find steady-state solution as seen on both plots after generation №500, while polynomial mutation (red line) results in poor [Ca^2+^]_nsr_ convergence. Dashed line in the both panels corresponds to input model concentration values.

### Elitism strategy

Genetic operators tend to spoil “good” solution; therefore, best organisms are passed to the next generation without any modifications^3,4^. We have found that in our GA implementation a large number of elite organisms is required for fast GA convergence. The SD dependence on the proportion of elite organisms to the whole population (Fig 6A) demonstrates that 6-10% of elite organisms is optimal.

**Figure 6.**
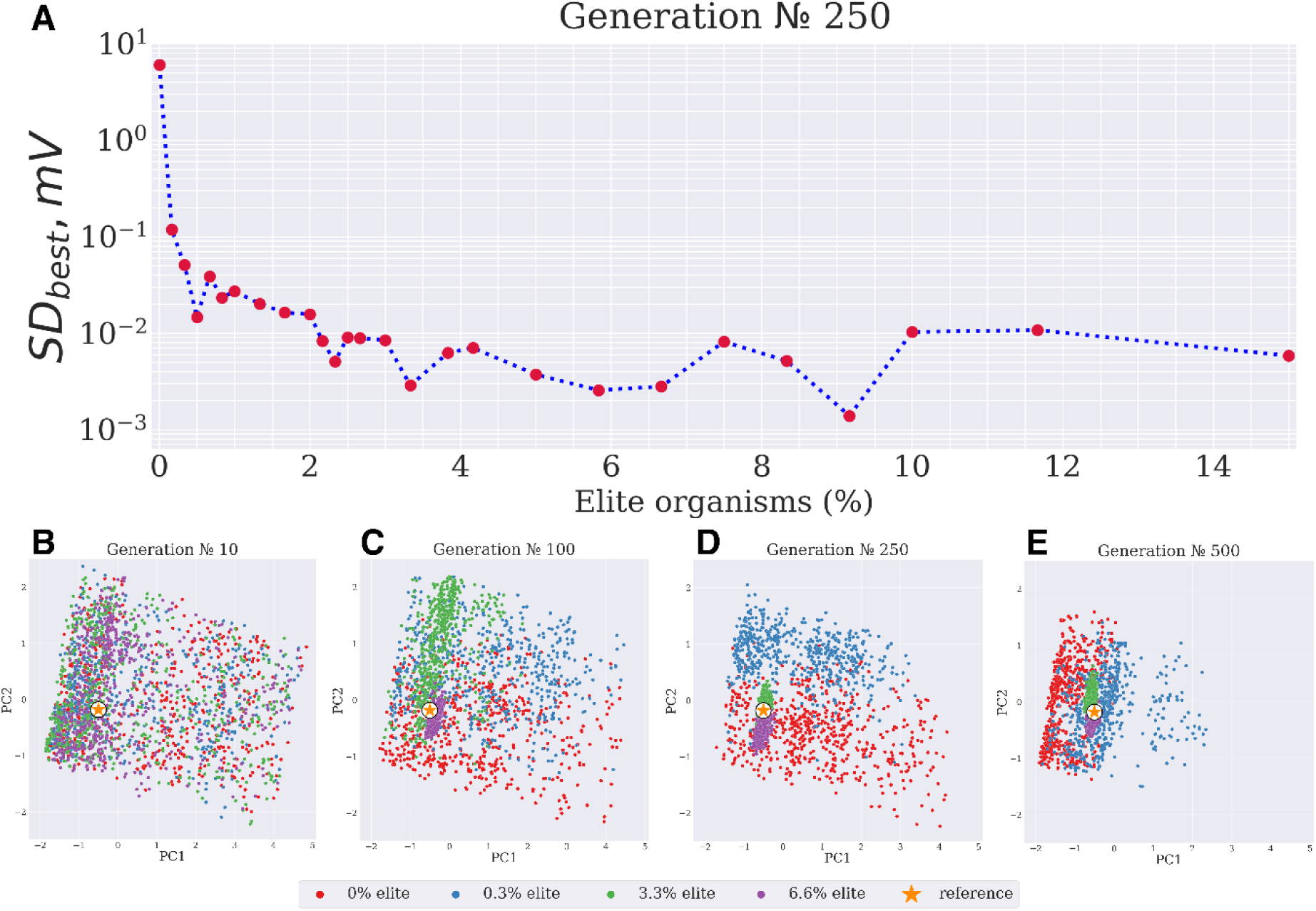
Modified elitism strategy. **(A)** SD dependence on the number of elite organisms. (**B**,**C**,**D**,**E**) Principal component analysis of convergence dependence on the number of elite organisms: 0% (red points), 0.3% (blue points), 3.3% (green points) and 6.6% (purple points). Higher number of elite organisms results in the faster clusterization around the input model parameters.

Principal component analysis (PCA) comparison of sample GA runs (Fig 6B-E) shows that in case of higher proportion of elite organisms (6.6%) solution tends to cluster around the precise solution at generation № 100, while a GA run with 3.3% of elite organisms requires at least twice the number of generations to converge the population to the same cluster size. As further explained in Fig 7 high proportion (6.6% or 3.3%) of elite organisms results in the fast reduction of the cluster size (Standard Distance of the population, Fig 7C). This reduction, in turn, allows to “fine-tune” the solution by the algorithm: mean cluster error (MCE) has a clear trend after the reduction of the cluster size (Fig 7B), while MCE for the low proportion of elite organisms (0% or 0.3%) follows random fluctuations.

**Figure 7.**
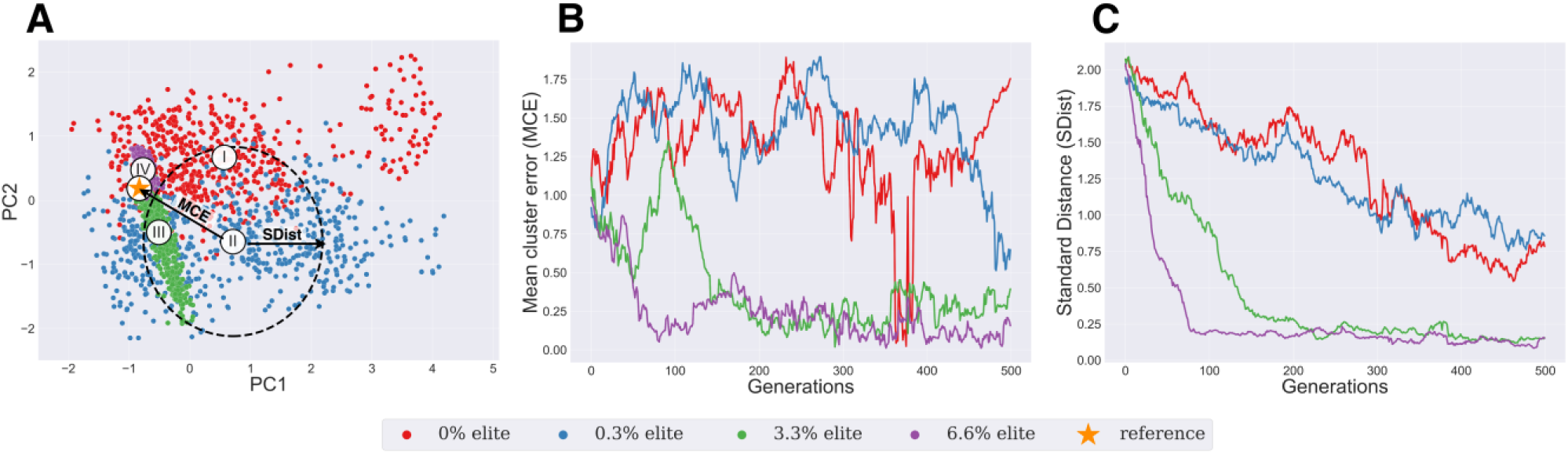
Clusters characteristics: Mean Cluster Error (MCE) and Standard Distance (SDist). **(A)** Mean cluster error (distance between the center of each cluster and reference value) and Standard Distance (plotted as a radius of dashed circle, measures the compactness of a distribution). Cluster mean centers are shown by numbers I (corresponding 0% of elite organism, red points), II (0.3% of elite organism, blue points), III (3.3% of elite organisms, green points), IV (6.6% of elite organisms, purple points). (**B**) MCE dependence on generation number for each cluster. Purple and green clusters rapidly shift to the exact solution area and remain there until the GA termination, while red and blue clusters don’t converge to the reference area. (**C**) SDist dependence on generation number for each cluster. Purple cluster size decreased approximately 8 times after a hundred of generations. Red and blue clusters size decreased 2.6 times after 500 generations.

### Final algorithm

In order to test if the new algorithm narrowed down the solution range, we have compared our algorithm with the original one^3^ (Fig 8). As explained above (“Intracellular concentrations” paragraph), it was computationally very expensive to achieve steady state with the original algorithm. Therefore, in the original algorithm each organism was paced for 50 stimulations at every PCL, *i.e.* quasi-steady-state was used instead. As shown in Fig 7 our algorithm determined I_K1_ conductivity with 7% precision, I_Kr_ – 5%, I_Na_ – 7%, I_CaL_ - 12% (vs. 39%, 29%, 30%, 203% maximal error, correspondingly, in the original algorithm). Membrane currents that did not have profound effects on the AP waveform were less precise (I_Ks_, I_to_, I_NCX_, I_NaK_ – 39%, 37%, 79%, 38% error, respectively), however, precision was still better then original algorithm (144%, 122%, 113%, 99%, correspondingly). Model parameters that didn’t affect AP waveform directly (RyR, SERCA and CAMKII) were also determined with an error less than 69%.

**Figure 8.**
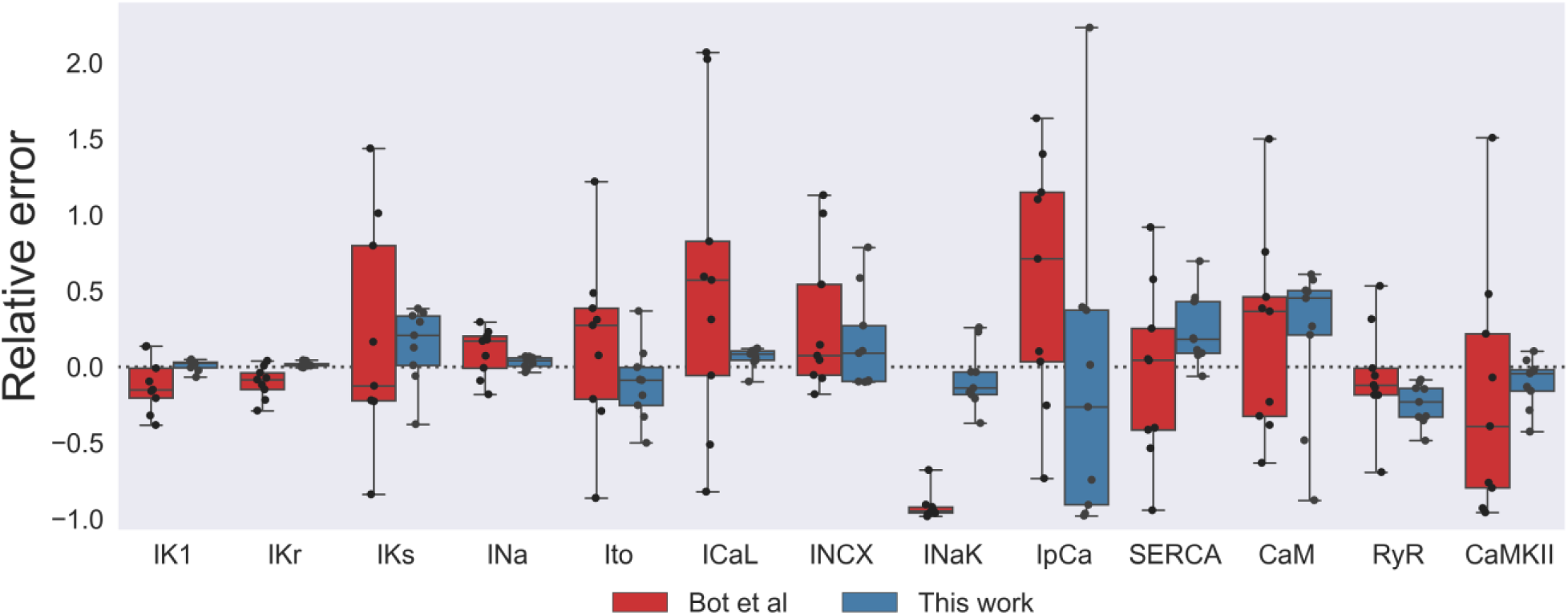
Comparison of presented algorithm precision to Bot et al. Parameter scaling by the presented algorithm (blue boxes) as compared to ^3^ (red boxes) after a long run (700 generations). Dashed line corresponds to input model parameter value.

### Input data requirements

In the results described above, input data was APs simulated at 7 different PCLs (see “Methods” section). However, in real experimental setting it is preferable to reduce the recording time (*i.e.* number of PCLs at which steady state AP waveform is recorded). We have tested GA performance, when input AP was simulated at limited number of pacing frequencies. Particular PCLs we used to compare algorithm performance are listed in Fig 9A. We observed that for some parameters (Ito, SERCA: S3E,J Fig) single AP waveform recorded at 1000 ms was enough (*i.e.* algorithm precision did not increase when more detailed restitution properties were used). Two extreme points on the restitution curve (217 ms and 2000 ms) were required to determine I_Kr_ conductivity with 5% precision (S3B Fig). The parameters most sensitive to the AP restitution are shown in Fig 9B-F. It is preferable to use full restitution curve (7 PCLs) for I_Ks_, I_Na_ and I_CaL_. However, 4 points on restitution curve still result in acceptable precision for every other parameter.

**Figure 9.**
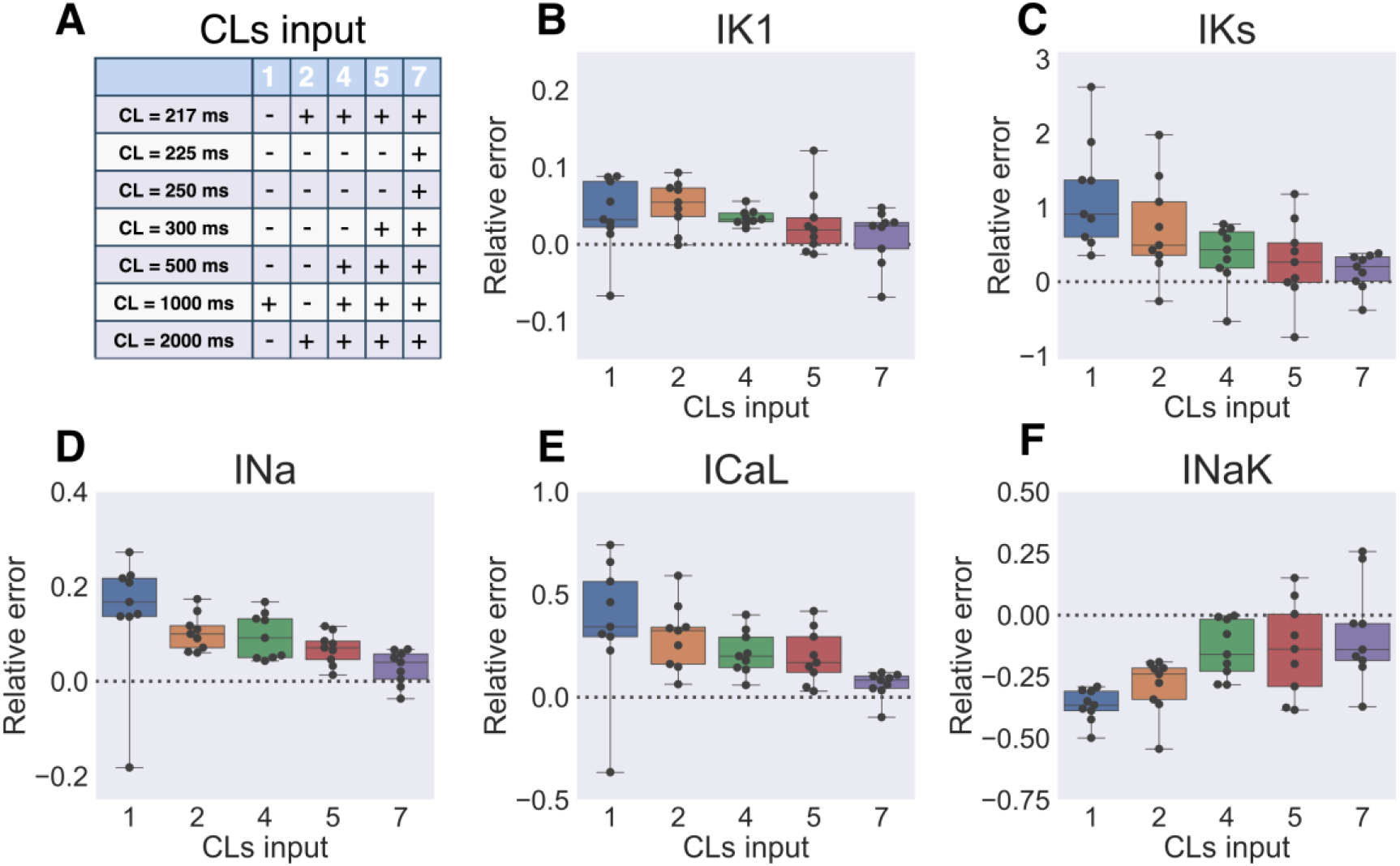
Solution sensitivity to the number of input baselines. (**A**) Input model AP was recorded at several PCLs listed in the table. Larger number of input baselines results in better algorithm performance. (**B-F**) Box-and-whiskers plots depict the parameters most sensitive to the changes in the number of input baselines (I_K1_, I_Ks_, I_Na_, I_CaL_, I_NaK_) at generation №700 of 8 GA runs. Dashed line corresponds to the input model parameter value.

The other inherent aspect of experimental data that might spoil algorithm performance is noise. To test how noisy data would affect the algorithm precision we have added Gaussian noise to the input simulated AP. Sample AP waveforms with different signal to noise ratio (SNR) are depicted on Fig 10E,F and S4 Fig. As expected, noise amplitude heavily affected the output parameters precision. As seen on Fig 10A-D and S5 Fig 20 dB SNR completely breaks down algorithm: for example, error is up to 100 % for I_Kr_ and up to 227% for I_Ks_ conductivities. 28 dB SNR, achievable in experimental setting gives better results: maximal error in 9 runs was 4% for I_Kr_, 14% for I_Na_, 31% for I_CaL_, 36% for I_NaK_. Although maximum error for I_Ks_ is still very high (99%), median value is much closer to the precise value (34% error) than in the case of high-amplitude noise.

**Figure 10.**
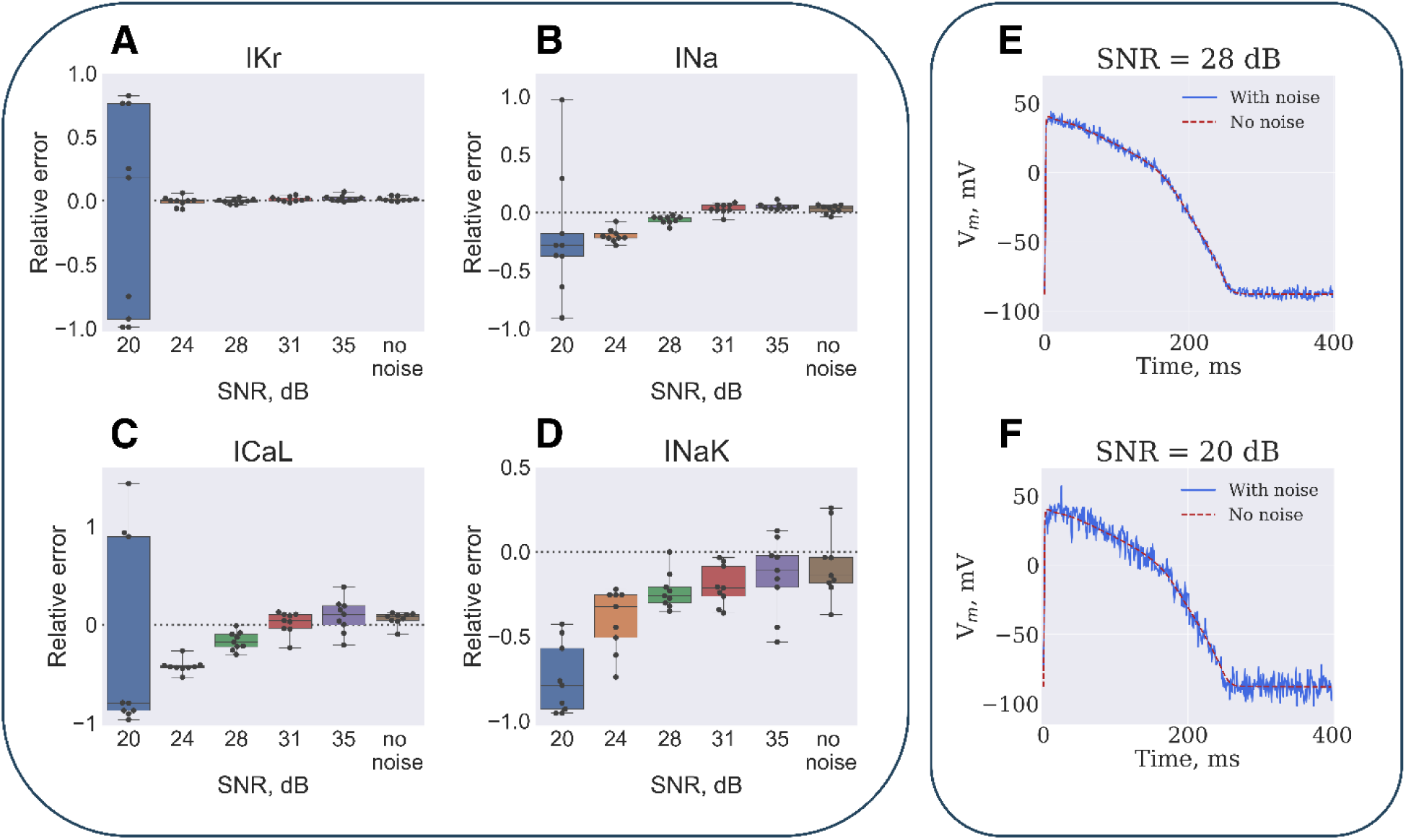
Solution sensitivity to the input baselines signal-to-noise ratio (SNR). The input model AP waveform was distorted by Gaussian noise. (**A**) Parameters dependence on the SNR is illustrated at generation № 700 of 9 GA runs. Dashed line depicts the input model parameter value. (**B**) Sample AP waveform at 28 dB and 20 dB SNR.

### Experimental data

GA was also tested with optical APs as input data (see Fig 1B and “Methods” section). Fig 11A compares output model with input data. The *Patient 2* model faithfully reproduced AP waveform dependence on the PCL, however, we observed slight deviations between model and experiment (up to 6 mV in membrane potential and up to 15 ms in AP duration). The membrane potential differences are mostly confined to depolarization phase and are probably due to photon scattering in optical mapping setting ^23^.

**Figure 11.**
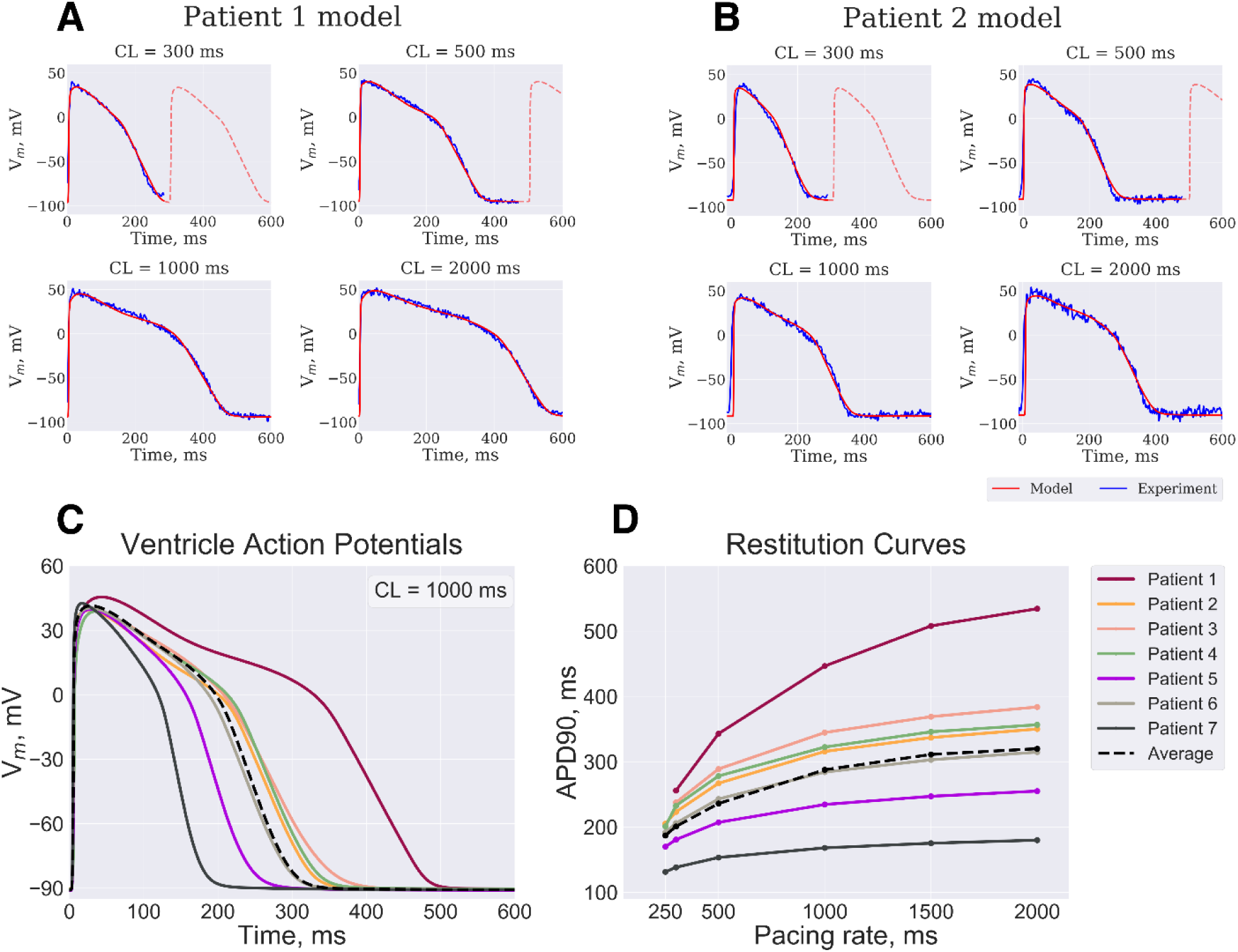
Algorithm verification. (**A**) Patient 2 (see text for details) model APs (red line) fitted by GA as compared to the optical APs (blue) recorded at CL of 2000 ms, 1000 ms, 500 ms and 300 ms. (**B**) Patient 1 model was rescaled using mRNA expression data and compared to experimental APs (see Fig. 1B). Personalized models based on mRNA expression data from 7 donor hearts ventricles demonstrate different AP waveform **(C)** and restitution properties **(D)**.

As described above (“Methods” section), in order to verify output parameter values the *Patient 2* model parameters were rescaled proportional to the difference of mRNA levels of expression between *Patient 1* and *Patient 2*. The resulting *Patient 1* model was compared to optical mapping data. *Patient 1* model still faithfully reproduced AP waveform at every PCL (Fig 11A), however, membrane potential and AP duration errors were somewhat more pronounced: up to 13 mV and 20 ms correspondingly. Using a similar approach we have reconstructed personalized models for other 5 patients, AP waveform and restitution curves are depicted in Fig 11C-D. The variability between the models is within physiological range^24^. *Patient 5* and *Patient 7* APD was too short (below 200 ms), which is explained by the fact that mRNA levels of expression had the most extreme deviations from median value in these patients. *Patient 5* was an outlier in terms of ATP1A1, ATP2A2, ATP1B3, ATP2C1, KCNJ5, KCNK3 genes. *Patient 7* was an outlier in terms of ATP1A1, ATP1B1, ATP1B4, CACNA1C, CACNA2D1, CACNA2D3, CACCB1, CACNB2, CALM1, CALM3, KCNH2, KCNJ11, KCNJ3, KCNK1, KCNK3, CAMK2B, CAMK2D genes (see also Fig 2). These deviations might indicate problems with heart preservation prior to tissue collection or undiagnosed heart diseases.

## Discussion

In this study, we introduce a genetic algorithm-based method for personalized for cardiac electrophysiology models reproducing patient-specific AP recordings. The advantages of the algorithm are summarized below.

1. Many different combinations of model parameters result in the same AP waveform. However, these solutions are mostly non-steady-state: AP waveform diverges from experimental in the long run. Steady-state requirement allowed us to substantially narrow down the parameters range. As shown above (Fig 8, S3 Fig), using AP waveform dependence on PCL as input data further increased algorithm precision and convergence to a unique solution.
2. Reaching steady state for every organism is computationally very expensive. To solve the problem, *parameter search* and *steady state search* are performed at the same time. Model gets closer to steady state after a short simulation, after that final model state is saved, the fitness function is evaluated, and parameters are modified by genetic operators. Next generation simulations start from new initial state bringing the model closer to steady state. This approach has two disadvantages: firstly, as demonstrated in Fig 3A-C steady state is initial-stated dependent; secondly, the slowest variables still require very long run (many generations) to approach steady state. Therefore, two slowest variables: intracellular sodium concentration and sarcoplasmic reticulum calcium load are treated as model parameters by the algorithm, *i.e.* mutation and crossover operator are applied to these variables as well. As shown in Fig 3D-E beneficial byproduct of this approach is that intracellular ionic concentrations are determined by the algorithm with a millimolar precision.
3. Traditional genetic algorithm tends to trap the whole population within a single local minimum (Fig 5C), after that global minimum might never be reached. We found it especially important in terms of slow variables: as shown in Fig 5E *Ca*^2+^_*NSR*_ did not converge to proper value even after 700 generations. This problem was solved by using Cauchy distribution for mutation operator: both moments (expected value and variance) are undefined for this “*pathological*” distribution, thus a number of outliers are always generated outside of the local minimum.
4. The convergence speed of the algorithm was improved by further modifications in the mutation operator and in the elitism strategy. We found that mutating each parameter with a fixed probability (referred to as “*point mutation*” above) resulted in coordinate descent (Fig 4B), which is very slow in 27-dimensional parameter space (parameters include intracellular concentrations at each PCL) and usually stopped convergence after about 100 generations (Fig 4C). Instead, we chose random direction in multidimensional parameter space and mutated parameter in this direction, resulting in better algorithm performance (Fig 4E, F). We also found that increasing the number of elite algorithms further improved convergence of the algorithm: 7% of elite organisms was optimal for our problem (Figs 6–7).
5. The resulting algorithm evaluates the most important model parameters (I_K1_, I_Kr_, I_Na_ and I_CaL_) with high precision: maximal error for these parameters was below 13% (Fig 8). However, low amplitude ionic currents and parameters affecting calcium transients were much less precise.
6. The algorithm sensitivity to noise in input data is acceptable: while 20 dB breaks the algorithm, SNR above 28 dB has minor effects on algorithm performance (Fig 10).

We did not experimentally measure ionic channels conductivities directly in this study, thus we could not directly verify algorithm precision in the experimental setting. However, we found that output model parameters are in accordance with mRNA expression profile (Figs 1B, 11). Another point should be discussed in this regard: while the algorithm is relatively stable, SNR requirements are quite strict. It was possible to achieve 28 dB level noise in *ex vivo* optical mapping experiment; however, it still required some post-processing: hum removal and ensemble averaging in particular. That probably implies that algorithm cannot be used in a clinical setting based on extracellular electrograms. Monophasic action potential recordings in clinical settings could solve this problem, because they reproduce the exact AP waveform. However, in some cases (for example, post heart transplant patients) ventricular biopsy could be justified, and tissue samples might be obtained from the patients’ heart. As shown in Figs 1B and 11 using genetic algorithm used in combination with gene expression profiling allows one to develop precise personalized model reproducing AP waveform as well as restitution properties.

## Limitations

The output conductivities of high-amplitude ionic currents were very precise; however, algorithm performance was much less accurate for important model parameters affecting calcium transients, RyR and SERCA in particular. Using multiparametric optical mapping^25^ as input data could probably further increase algorithm precision, however, this question is beyond the scope of the current study.

We used a rough approach for mRNA-based personalized models: ionic channel conductivity was taken proportional to a single gene level of expression. Mostly we used the pore-forming protein for this purpose. More precise approach would require one to account for auxiliary subunits affecting ionic channels voltage-dependence.

## Acknowledgments

The research was supported by Russian Foundation for Basic Research grants 18-07-01480, 19-29-04111, 18-00-01524 and Leducq Foundation (project RHYTHM).

