## Supplemental materials for "Genetic algorithm-based personalized models of human cardiac action potential"

### SUPPLEMENTARY MATERIALS

**Tissue simulation.**

The algorithm developed in the current study was optimized to reconstruct model parameters using optical mapping action potential (AP) recordings in the cardiac tissue. Intercellular interactions affect AP waveform; therefore, 1D tissue was simulated to evaluate the fitness function. For each pacing cycle length (PCL) the following system of ordinary differential equations was solved:

$$\frac{dV_m}{dt} = -\frac{1}{C_m}\sum_i I_i + I_{gap} = -\frac{1}{C_m}\sum_i I_i + g_{gap}(V_{m-1} - 2V_m + V_{m+1}),$$

where $V$ is the membrane potential, $C_m$ – cell membrane capacitance, $I_i$ – transmembrane ionic currents, $I_{gap}$ – junctional current, $g_{gap}$ – gap junctions conductivity, $V_{m+1}$ and $V_{m-1}$ are downstream and upstream cells, the size of a cell taken to be *100 μm*.

We have found that in case of relatively low conduction velocity (CV) value of *27 cm/s*, 30 cells-long (*3 mm*) tissue is adequate to minimize the boundary effects on the AP waveform (S1A Fig). On the other hand, 100-cells (*1 cm*) long tissue simulations demonstrate that within physiological range (*20-100 cm/s*) exact CV value does not affect exact AP waveform (S1B Fig). Therefore, a combination of *3 mm* long tissue size and *27 cm/s* CV was used in all genetic algorithm (GA) runs in this study.

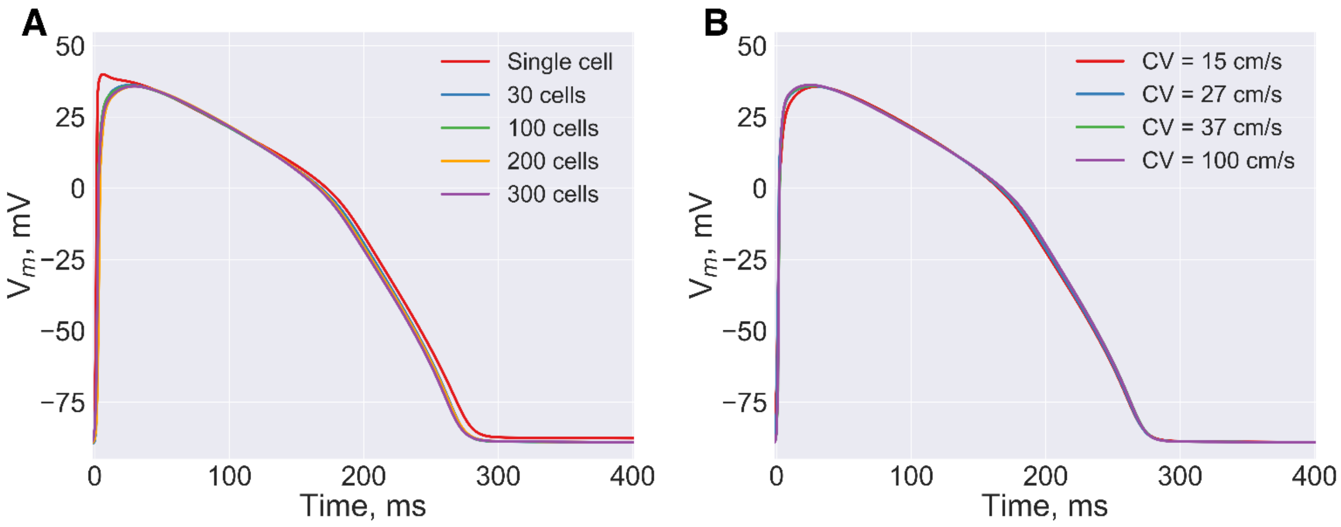


**S1 Fig. Tissue effects.**
(A) Comparison of a single cell AP waveform (red line) and AP waveforms recorded from a central cell of a 1D string of cells, tissue size was varied, CV was ≈*27 cm/s* in tissue simulations. (B) Comparison of AP waveforms recorded from a central cell of a 100-cells long (*1 cm*) string of cells with variable gap junctions conductivity.

#### Cauchy mutation.

As discussed in the main text of the article, Cauchy mutation operator allows the algorithm to exit the local minima, which we found to be most important to find appropriate steady state values of slow variables. Fig 5D,E of the main text compares two GA runs to demonstrate that steady state was found in case of Cauchy, but not Polynomial mutation. S2 Fig demonstrates slow variables dynamics averaged over 9 GA runs, demonstrating that Cauchy mutations results in better convergence of slow variables.

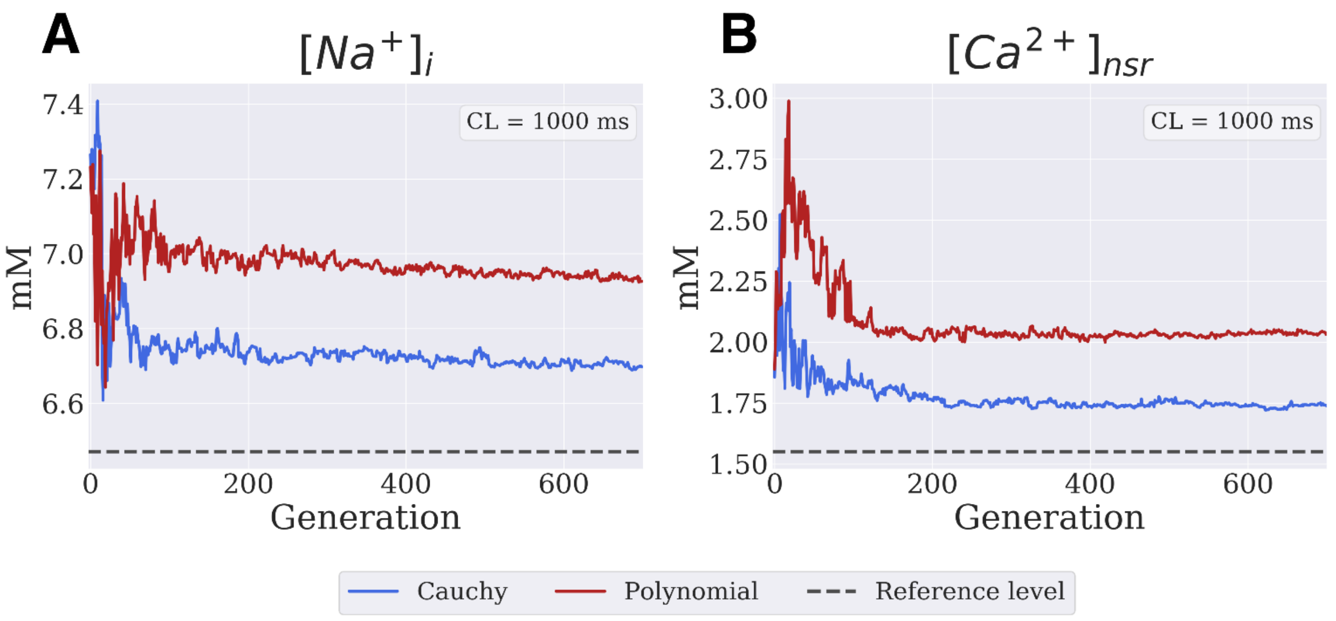


**S2 Fig. Dynamics of $[Na^+]_i$ and $[Ca^{2+}]_{nsr}$ concentration.**
Best organism intracellular $[Na^+]_i$ and $[Ca^{2+}]_{nsr}$ concentration averaged over 9 GA runs plotted against generation number. Dashed line in both panels corresponds to input model concentration values.

#### Input data requirements.

The algorithm developed in this study is based on AP waveform dependence on PCL. Since it is usually preferable to reduce restitution protocol time in experimental setting (*i.e.* the number of PCLs at which steady state AP waveform is recorded), we have tested how limited number of input baselines affect algorithm performance. S3 Fig summarize output parameters sensitivity to the number of input AP waveform recordings. Particular PCLs we used to test output parameters precision are listed in S1 Table below.

**S1 Table. PCLs input.**

| | 1 | 2 | 4 | 5 | 7 |
|---|---|---|---|---|---|
| **CL = 217 ms** | - | + | + | + | + |
| **CL = 225 ms** | - | - | - | - | + |
| **CL = 250 ms** | - | - | - | - | + |
| **CL = 300 ms** | - | - | - | + | + |
| **CL = 500 ms** | - | - | + | + | + |
| **CL = 1000 ms** | + | - | + | + | + |
| **CL = 2000 ms** | - | + | + | + | + |

Input model AP was recorded at several PCLs listed in the table. In order to test how restitution protocol level of detail affects algorithm precision, we have varied number of input APs ("+" marks PCLs that were used as algorithm input in a particular GA run).

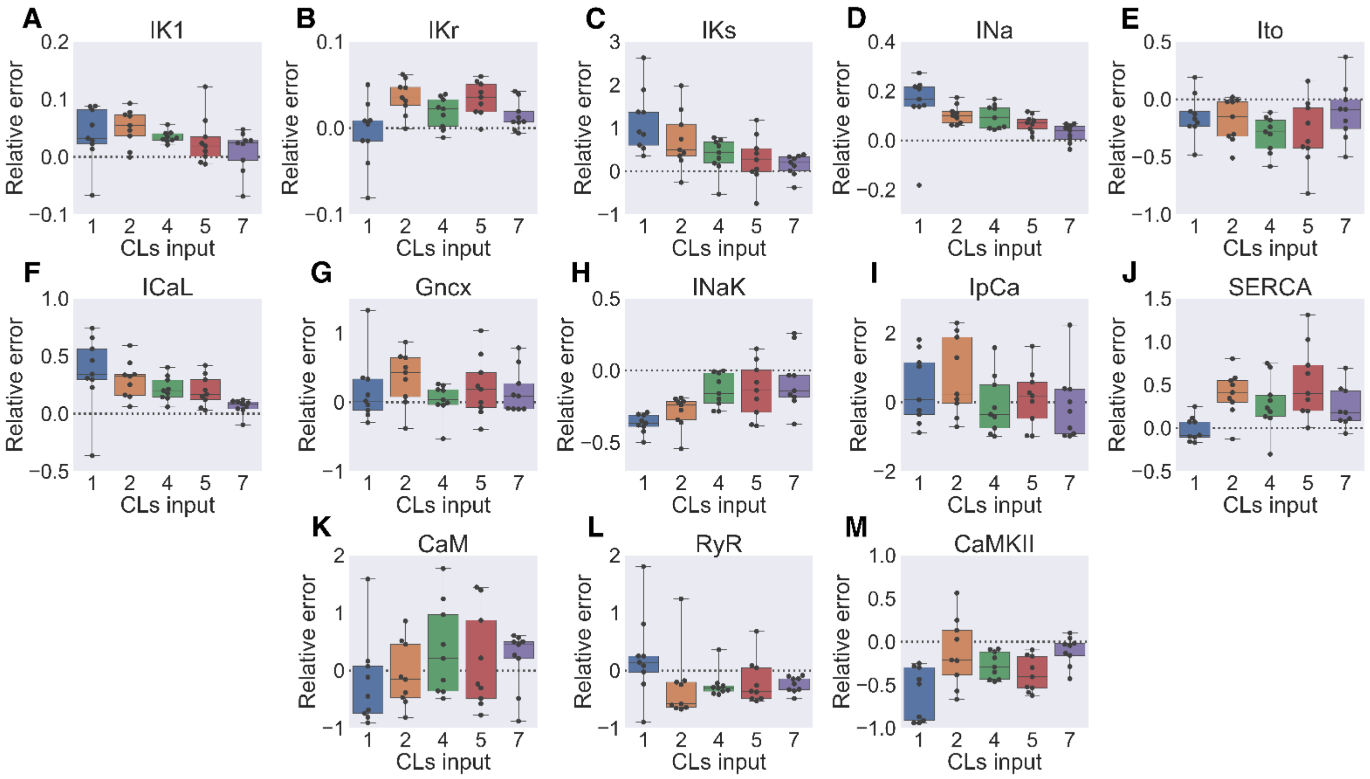


**S3 Fig. Solution sensitivity to the number of input baselines.**
(A-M) Box-and-whiskers plots depict the model parameters sensitivity to the number of input AP baselines. Input AP was simulated at several PCLs listed in the S1 Table. Dashed line corresponds to the input model parameter value.

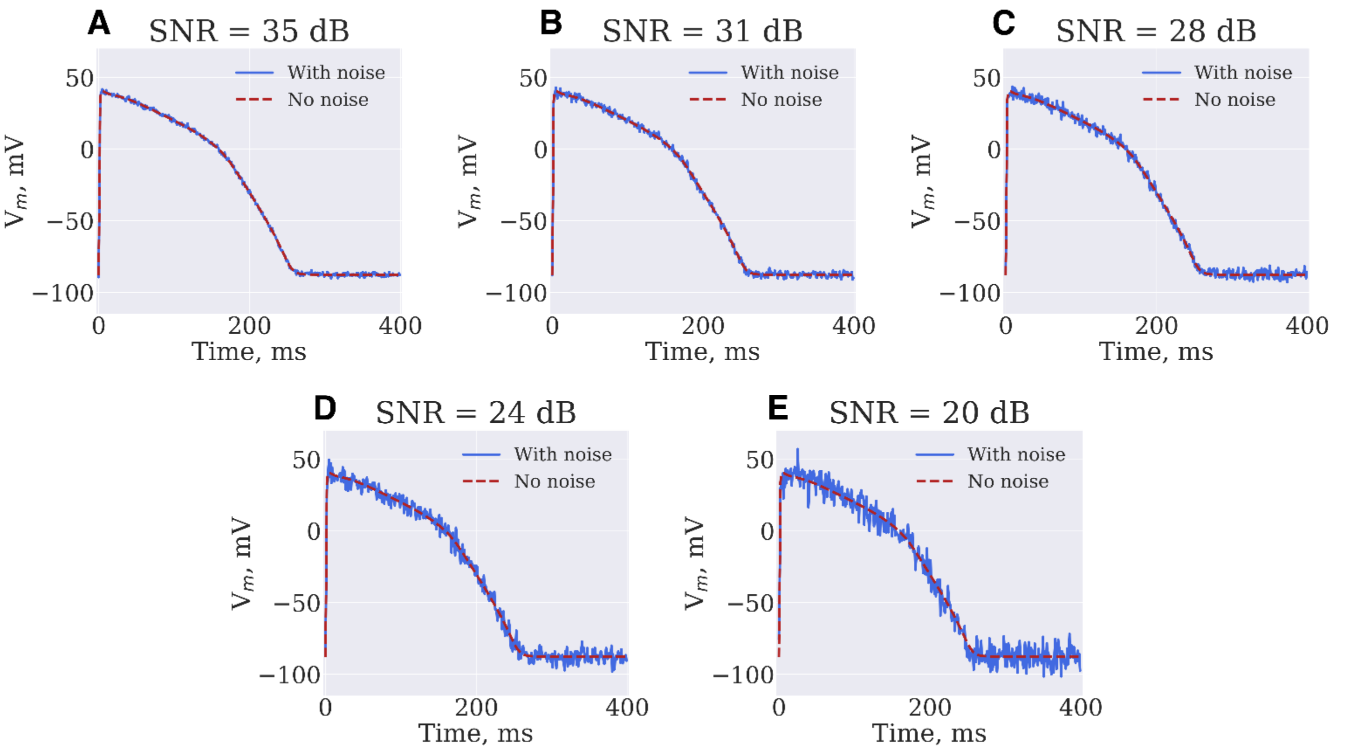


**S4 Fig. Input baselines signal-to-noise ratio.**
(A-E) APs waveforms (blue curves) for the different SNR values: 35 dB, 31 dB, 28 dB, 24 dB, 20 dB. Red dashed lines correspond to precise signal with CL = 1000 ms.

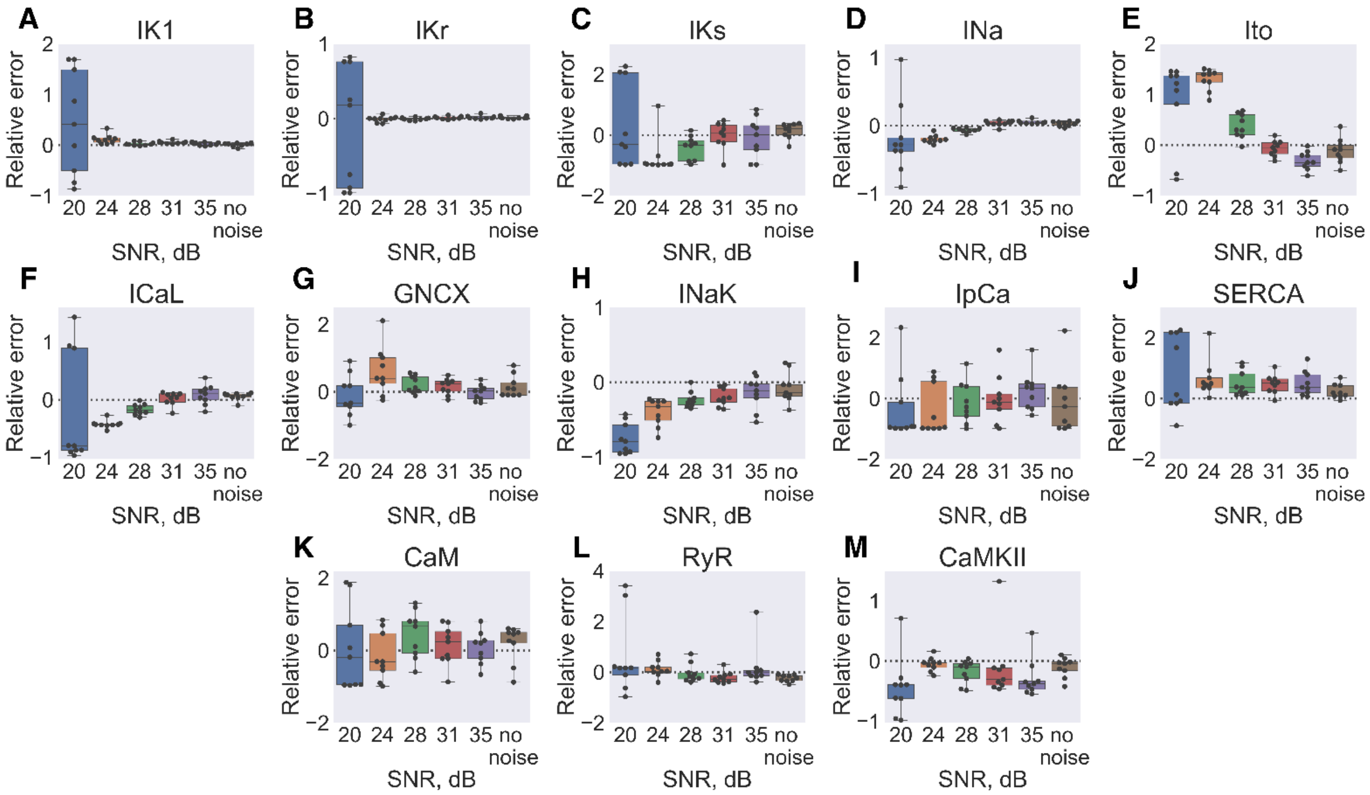


**S5 Fig. Parameters dependence on the SNR.**

(A-M) Optimized model parameters distribution depending on the SNR of input APs. Dashed line depicts input model parameter value.

In order to test algorithm sensitivity to input AP signal-to-noise ratio (SNR), simulated input APs waveforms were distorted by Gaussian noise (S4 Fig). S5 Fig summarize output parameters sensitivity to SNR. To verify the assumption that experimental noise follows normal distribution in the optical mapping experiment, we have calculated the difference between input optical action potentials (OAP) and their corresponding GA output APs:

$$\Delta V_i = \left| V_{\exp}(i) - V_{\mathrm{mod}}(i) \right|, i = t_{start}, \ldots, t_{end} ,$$

where $V_{exp}$ is experimental membrane potential, $V_{\mathrm{mod}}$ corresponds to model membrane potential, thus GA output AP was supposed to correspond to undistorted experimental AP. Experimental OAP were preprocessed with narrow band stop IIR Butterworth filter to remove 60 Hz-hum, furthermore, depolarization phase was excluded from the set (to exclude photon scattering effects from the consideration). As demonstrated by histogram and probability plot on S6 Fig experimental noise is indeed close to normal distribution.

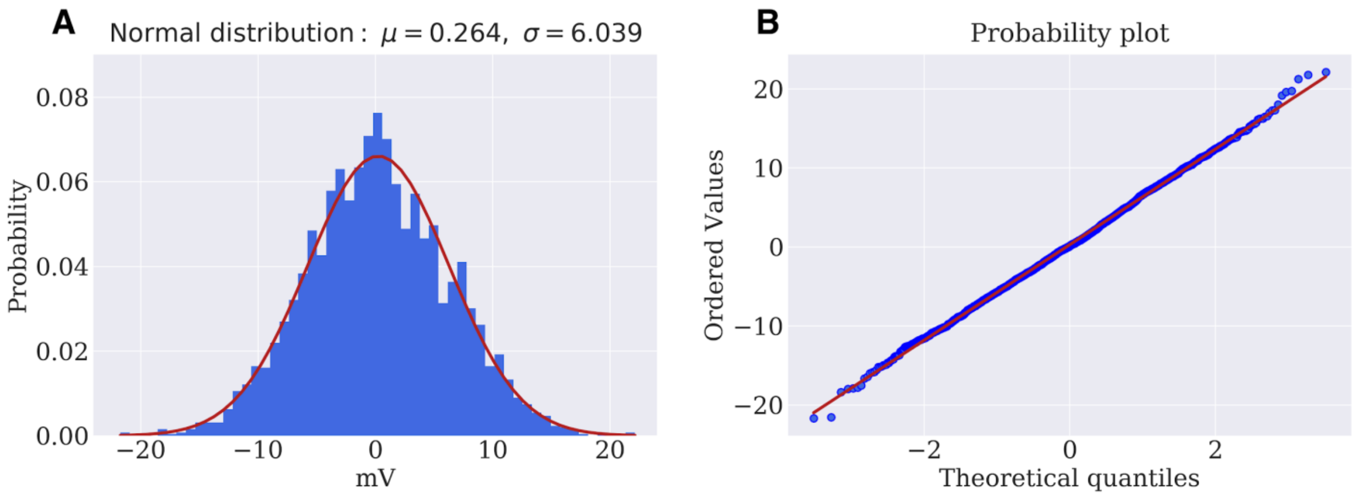


**S6 Fig. Gaussian noise.**
(A) Experimental noise is reproduced the normal distribution with mean = 0.264 mV, and standard deviation = 6.039 mV.
(B) Corresponding probability plot: quantiles of experimental noise amplitude distribution (blue) are plotted against quantiles of a theoretical normal distribution (red line).

#### Source code

The current version of the presented Genetic Algorithm is available to use. Link: https://github.com/humanphysiologylab/Genetic-Algorithm.
